# How to enviromically predict breeding genotypes as if they were commercial cultivars?

**DOI:** 10.1101/2025.05.28.656616

**Authors:** Rafael Tassinari Resende

## Abstract

Bridging the gap between how breeding genotypes perform in trials and how they might fare as commercial varieties or cultivars remains one of the enduring challenges in plant breeding, further compounded by the complexities of genotype-by-environment (G×E) interactions. This study implemented and evaluated an unstructured – ‘uns’ – bivariate enviromic reaction-norm model capable of predicting breeding genotypes ‘as cultivars’ by integrating genomic and environmental data. All model components were developed from scratch in R, ensuring full methodological transparency and reproducibility. The model jointly estimated genetic and residual variance-covariance matrices for experimental trials and commercial stands using a derivative-free restricted maximum likelihood (DF-REML) approach with the BOBYQA algorithm. Results demonstrated that employing a SNP-based genomic relationship matrix yielded biologically consistent estimates and enabled the construction of spatial recommendation maps for genotype deployment across diverse environments (e.g., in fields established by farmers or growers). These findings underscore the importance of modeling experimental *versus* commercial performance as distinct but genetically correlated traits, while also demonstrating the feasibility of implementing complex enviromic models without reliance on specialized software. This approach offers methodological advances for predictive breeding, supporting more efficient selection and reducing reliance on extensive multi-environment trials.

## 1 Introduction

Figuring out—or better yet, forecasting—how breeding genotypes under evaluation would perform as commercial cultivars remains a key challenge for the plant breeder. Genotypes are tested in breeding trials with the expectation that some will eventually be released as cultivars (Podlich et al., 2001; Bernardo, 2010). However, multi-environment trials (METs), despite providing structured phenotypic evaluations, often fail to encompass the environmental diversity encountered in actual farming conditions (Resende et al., 2021; Crossa et al., 2021). Genomic selection, leveraging dense molecular markers, improves prediction within observed environments but struggles to extrapolate to untested settings due to complex genotype-by-environment (G×E) interactions (Crossa et al., 2021; Heslot et al., 2015). Cooper et al. (2021) and Cruz et al. (2025) emphasize that METs frequently misrepresent the true target population of environments (TPE), highlighting the need to develop integrative predictive frameworks. Such integrative strategies are increasingly promoted in modern breeding, with recent frameworks emphasizing end-to-end predictive modeling to accelerate cultivar development by effectively linking early-stage evaluations with commercial deployment (Bhosale et al., 2025).

Experimental trials and commercial fields differ substantially in design, management, and environmental exposure. Trials are typically managed under optimal or uniform conditions, with replicated plots and controlled inputs, minimizing residual variation (Aparicio et al., 2024). Conversely, commercial fields exhibit substantial heterogeneity due to differences in soil types, climate variability, management practices, and biotic stresses (Messina et al., 2018; Milad et al., 2023). Thus, even when the same nominal trait (e.g., yield in kg/ha) is measured across both experimental and commercial settings, the underlying environmental variance structure renders these effectively different traits (Laidig et al., 2008; Cruz et al., 2025). Cooper et al. (2021) point out that breeding programs often overlook explicit mapping between METs and TPEs, risking overestimation of expected performance in real-world conditions. This justifies modeling experimental and stand trials as distinct but correlated traits, enabling the statistical model to capture their shared genetic basis while accounting for environmental divergence.

Reaction-norm models offer a formal framework to integrate genomic and environmental information, explicitly modeling G×E interactions (Jarquin et al., 2014; Lopez-Cruz et al., 2020). These models parameterize genotype-specific responses to envirotypic covariates (ECs) through random regressions, capturing both plasticity and stability across environments (Heslot et al., 2014; Monteverde et al., 2023). In the present approach, each genotype is modeled with a random intercept and random slopes on multiple ECs, separately for experimental and stand trial contexts. The genetic covariance structure is represented by a block-diagonal matrix, (*G*_0_ ⊗ *A*), where *A* is the genetic relationship matrix (pedigree- or SNP-based) and *G*_0_ is a 2 × 2 matrix capturing genetic variances and covariances across the two trait contexts. Separate residual variances are also estimated for experimental and stand trials (*R*_0_), accounting for distinct sources of environmental noise (Trevisan et al., 2025). This structure facilitates the emergence of a shared genetic correlation via joint reactions to ECs, enabling more accurate prediction of genotype-specific performance in both trial and commercial settings.

Accordingly, we conceptualize experimental and commercial stand performance as distinct yet genetically correlated traits. The genetic covariance matrix *G*_0_ captures the extent of shared genetic control, while *R*_0_ models environment-specific residual variability (Resende et al., 2021). This acknowledges that genotypic rankings may shift between trial and commercial contexts, a phenomenon well documented in G×E studies (Eskridge and Mumm, 1992; Crossa et al., 2021). Tilhou & Casler (2022) advocate for treating disparate evaluation contexts as “distinct but correlated” traits, enabling more nuanced selection decisions. The estimated genetic correlation indicates how well trial performance predicts commercial outcomes: high values suggest predictability, low values imply re-ranking. This bi-trait framework leverages genetic correlations to improve breeding value estimation for deployment.

The objective of this study is to implement and evaluate a bivariate (or bi-trait) enviromic reaction-norm model using a derivative-free restricted maximum likelihood (DF-REML) approach, specifically the “Bound Optimization BY Quadratic Approximation” (BOBYQA) algorithm (Powell, 2009). This model jointly estimates the genetic covariance matrix *G*_0_ and residual variance matrix *R*_0_ for experimental and commercial stands, allowing the prediction of commercial performance for breeding genotypes not yet tested under field conditions. Best linear unbiased predictions (BLUPs) for commercial stand yield are derived using each genotype’s experimental observations, its genomic relationships, and the associated ECs (Resende et al., 2024). Demonstrating accurate *in silico* prediction of commercial-like performance would provide a pathway for reducing the reliance on exhaustive METs, thereby expediting cultivar development pipelines (Gevartosky et al., 2023; Resende et al., 2024). This work applies the enviromic modeling paradigm to bridge the gap between experimental evaluation and practical deployment in modern breeding programs (Resende et al., 2021; Costa-Neto et al., 2021a).

## 2 Material and Methods

### 2.1 Genotypic, Phenotypic, and Envirotypic Data

The dataset utilized in this study comprised simulated phenotypic records, genotypic marker data, and envirotypic covariates (ECs) representing multiple locations and trial types. Two distinct types of observations were considered: *experimental trials* (EXP), corresponding to multi-environment trials (METs) conducted under controlled breeding conditions, and *commercial stands* (CST), representing real-world farming environments where genotypes are cultivated as commercial cultivars. Each observation was characterized by the following variables:

- **Geographic coordinates:** longitude and latitude of each trial location;
- **Location identifier (LOC):** code indicating the trial site;
- **Field type (FTP):** either EXP (experimental) or CST (commercial stand);
- **Genotype code (GCD):** identifier for each genotype, including both candidate genotypes (prefixed by ‘GEN’) and commercial cultivars (‘CUL’);
- **Phenotypic response (Y):** observed trait value (e.g., grain yield in ton/ha); for EXP, values are BLUEs or deregressed BLUPs by genotype and trial, while for CST, they are genotype-wise field means;
- **Envirotypic covariates (EC1 to EC5):** standardized variables representing local descriptors of each environment. Further details on their conceptualization and derivation are in Costa-Neto et al. (2021a) and Resende et al. (2024).

The dataset was simulated to facilitate clear visualization of each genotype and envirotypic covariate (EC), within a challenging scenario designed to stress-test the model implementation. Despite its modest size, the chosen optimizer — BOBYQA — is adept at efficiently handling large-scale problems with high-dimensional parameter spaces. This choice aligns with its application in packages like `lme4`, where derivative-free optimizers such as BOBYQA are routinely employed for mixed model estimation, even in extensive datasets (Bates et al., 2015). Consequently, the methodological pipeline remains scalable and applicable to real-world breeding data involving more complex structures.

The data structure, including phenotypic responses across trial locations, is illustrated in Figure 1. To understand the simulation process adopted here, readers may consult Resende et al. (2021), which presents a framework on enviromic-assisted selection. Specifically, a two-step algorithm is described: first, generating and characterizing the land area and envirotypic data; second, simulating trials, genotypes, and phenotypes by incorporating genotypic values, envirotypic interactions, trial effects, and residual variance.

**Figure 1.**
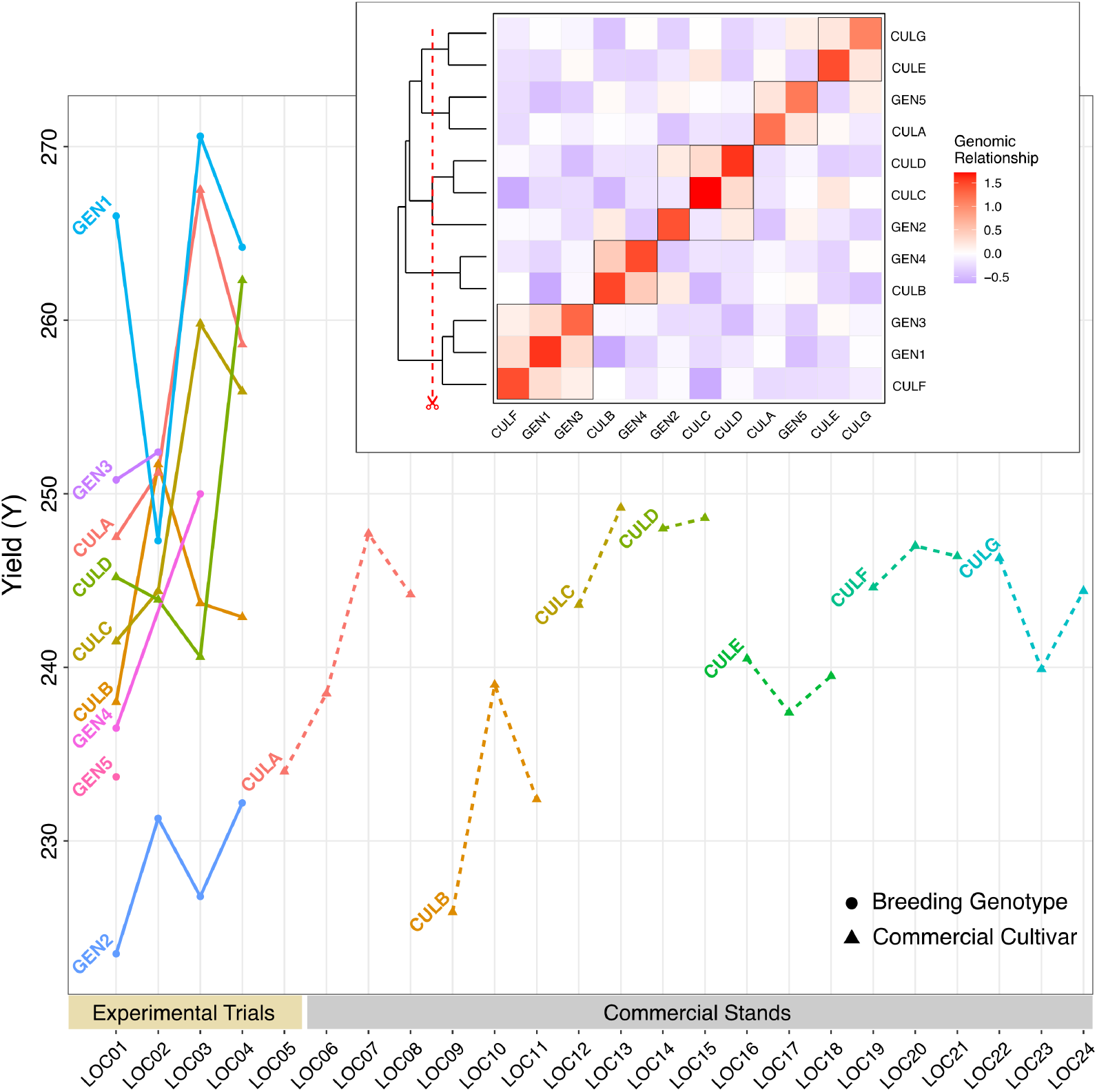
Background panel: phenotypic performance of genotypes across experimental trials (LOC01–LOC05) and commercial stands (LOC06–LOC24), with breeding genotypes represented by circles and commercial cultivars by triangles. Inset panel: clustered genomic relationship matrix (**A**), showing realized additive genetic similarities among genotypes.

**Figure 2.**
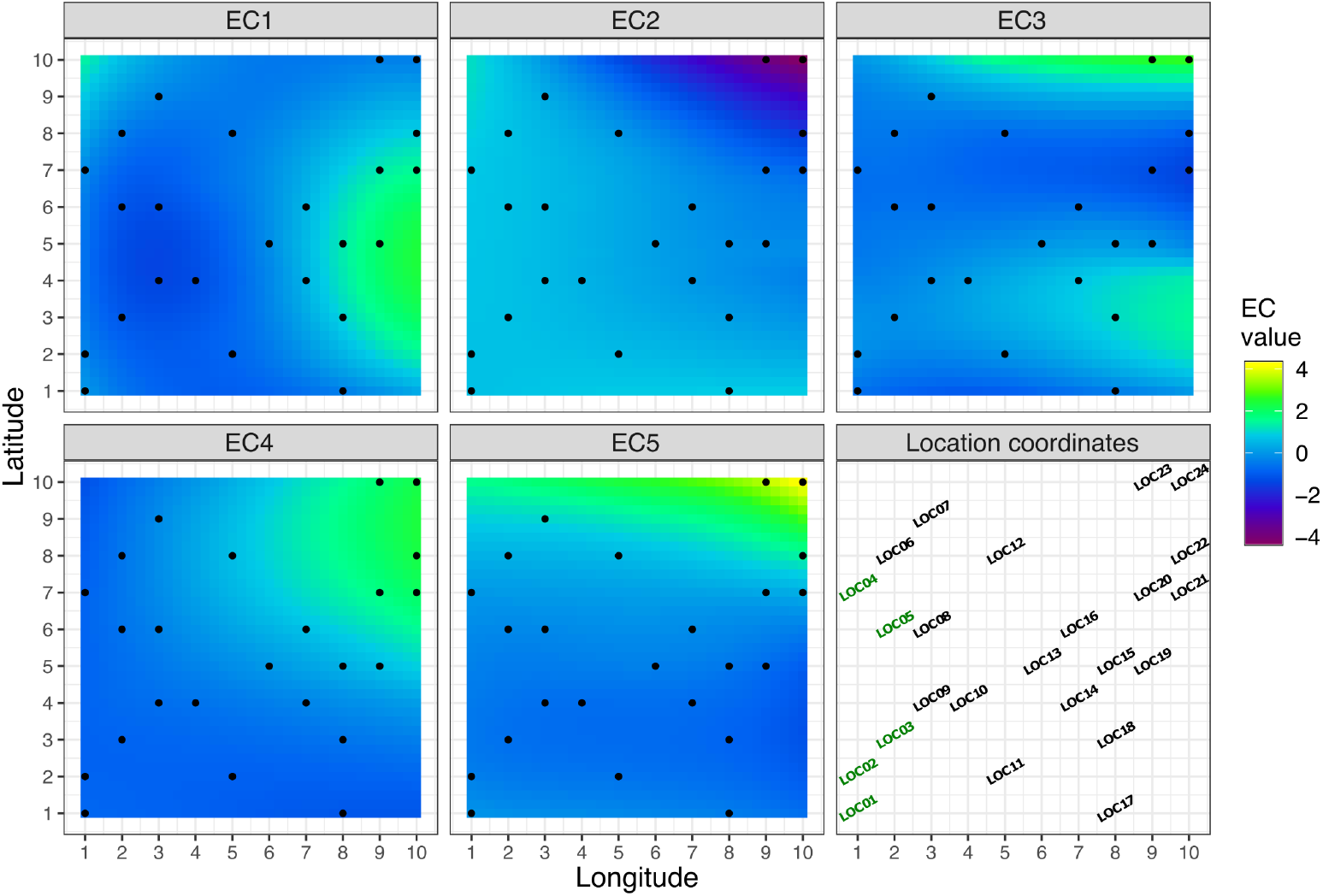
Spatial interpolation of envirotypic covariates (EC1 to EC5) across the study area, with observed locations indicated by black points. The bottom right panel shows the coordinates and labels of all locations (LOC01 to LOC24). Color gradient represents the EC values from low (blue) to high (yellow). Locations LOC01 to LOC05 (in green) correspond to environments where breeding experiments were conducted, whereas LOC06 to LOC24 represent commercial field environments.

**Figure 3.**
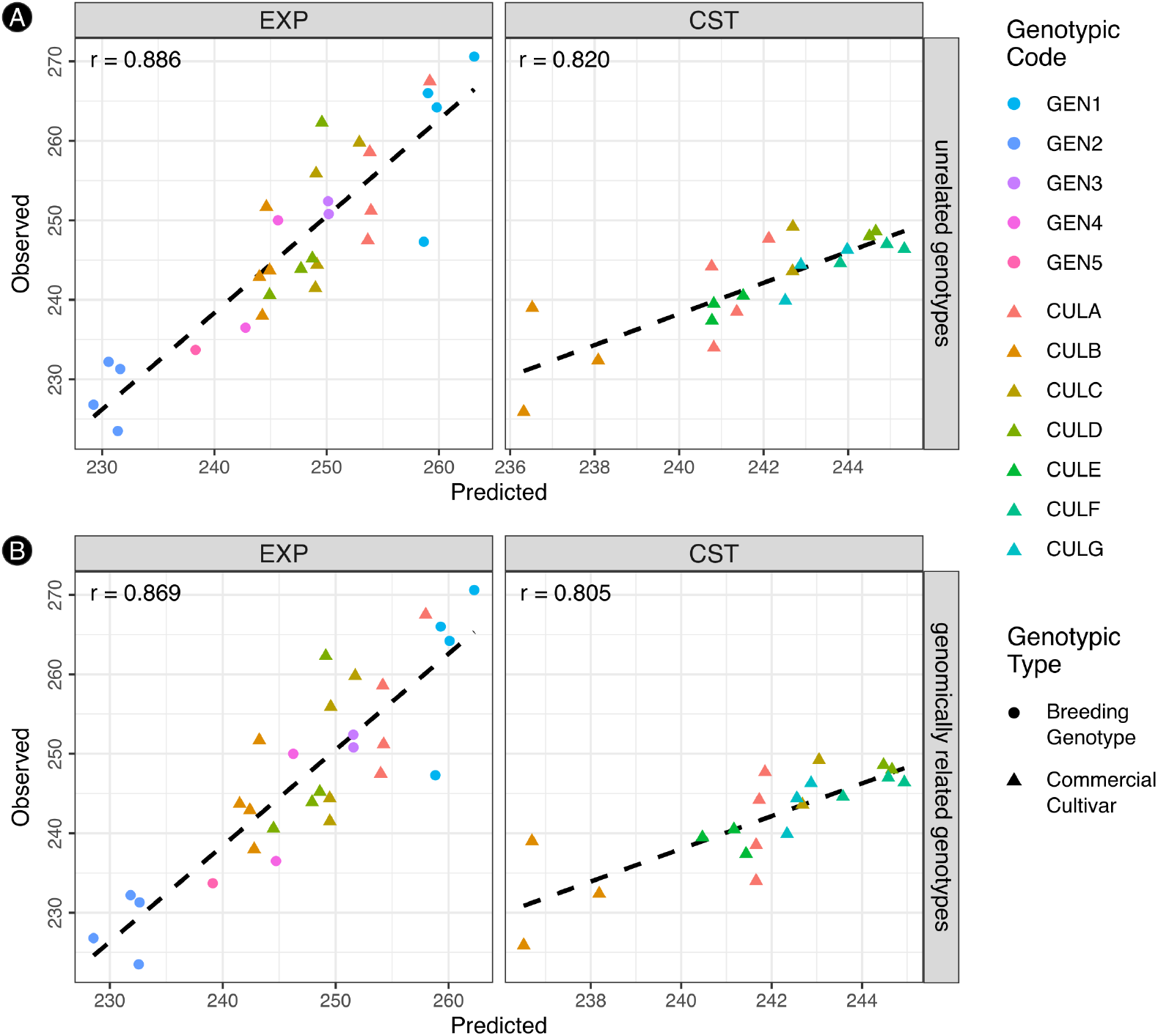
Predicted vs. observed phenotypic values for experimental trials (EXP) and commercial stands (CST), under two modeling scenarios. (**A**) Genotypes considered as *unrelated* (**A** = **I**_**12**_). (**B**) Genotypes considered as *genomically related* (SNP-based **A**). Circles represent breeding genotypes, and triangles represent commercial cultivars. Pearson correlation coefficients (*r*) between predicted and observed values are displayed within each panel. Note that these are in-sample fits, without cross-validation, thus providing an assessment of goodness of fit.

Genomic data, comprising fifty Single Nucleotide Polymorphism (SNP) markers coded additively as 0, 1, or 2, were used to compute the additive genomic relationship matrix (**A**), representing genetic similarities among genotypes. **A** was calculated using the `AGHmatrix` package (Amadeu et al., 2023), which centers **M** by subtracting twice the allele frequency; the centered matrix then yields:

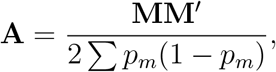

where *p*_*m*_ is the allele frequency at the *m*-th marker. This method estimates realized additive genetic similarity, accounting for linkage disequilibrium. The matrix **A** (also known as **G** matrix), was used in the bivariate mixed model to represent genetic covariance, enabling joint modeling of genotypic effects across both trial types (EXP and CST) and supporting enviromic prediction. The genomic **A** matrix appears in Figure 1 (*inset panel*).

### 2.2 Bivariate Mixed Model Implementation and Computational Details

This parameter estimation approach aligns conceptually with the framework presented by Resende et al. (2025) and Trevisan et al. (2025), who emphasize the construction of enviromic matrices within the mixed models basis. However, while their work focuses on theoretical formalization in a univariate setting, I extended the methodology to a bivariate model with full DF-REML implementation from scratch.

Moreover, this approach complements recent advances in enviromic modeling, such as the integration of engineered enviromic markers through ensemble modeling techniques proposed by Resende et al. (2025), illustrating the versatility and practical relevance of mixed model frameworks in modern plant breeding.

The fitted values obtained from the model were compared to the observed phenotypic responses for each genotype and environment combination to assess model fit. This comparison was visualized through scatter plots and Pearson correlation coefficients between predicted and observed values across field types.

#### 2.2.1 Construction of Fixed and Random Effects Design Matrices and Mixed Model Equations

The bivariate mixed model implemented in this study can be expressed, for each observation *i* = 1, …, *N*, as:

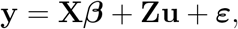

where **y** ∈ ℝ^*N*^ is the vector of phenotypic observations, ***β*** ∈ ℝ^2^ is the vector of fixed intercepts corresponding to each field type (experimental – EXP, and commercial stand – CST), **u** ∈ ℝ^2(1+*K*)*G*^ is the vector of genotype-specific random effects (intercepts and slopes for ECs), and ***ε*** ~ 𝒩 (**0, R**) is the residual vector with heterogeneous variances by field type.

The fixed-effects design matrix **X** ∈ ℝ^*N ×*2^ was constructed to model two intercepts, one for each field type *f*. This parameterization allows the model to capture distinct baseline performances across experimental and commercial stand environments. The random-effects design matrix **Z** ∈ ℝ^*N ×*2(1+*K*)*G*^ was constructed to accommodate, for each genotype *g* = 1, …, *G* and field type *f* = 0 (EXP) or 1 (CST), both a random intercept and *K* = 5 random slopes associated with ECs (*k* = 1, …, 5). Consequently, each genotype contributes 2(1 + *K*) random effects, accounting for its specific response in each field type.

The systematic procedure for constructing **Z** involved the following steps: **(1)** For each observation *i*, the corresponding genotype index *g* and field type index *f* were determined; **(2)** The **base column position** within **Z** was calculated as: base = (*g* − 1) × 2(1 + *K*) + *f* × (1 + *K*). This ensures that random effects for each genotype and field type occupy distinct, non-overlapping blocks of columns; **3)** At this base position, the first element was set to 1, representing the random intercept for genotype *g* under field type *f*; **(4)** The subsequent *K* elements were filled with the corresponding envirotypic covariate values from the dataset, representing the random slopes for each envirotypic covariate.

Thus, for each observation, exactly (1 + *K*) elements of a specific block within **Z** were filled: the intercept followed by the covariate values, with all other elements remaining zero, resulting in a sparse, structured design matrix.

Under the assumptions of the mixed model, the random effects follow **u** ~ 𝒩 (**0, K**), where **K** = **A**⊗(**G**_0_⊗**I**_1+*K*_) represents the covariance matrix of the random effects, with **A** representing either the identity matrix of order 12 (**I**_12_, ie., all 12 genotypes are unrelated) or the SNP-based genomic relationship matrix (simply referred to as **A**); **G**_0_ the genetic variance-covariance matrix between field types; and **I**_1+*K*_ an identity matrix accounting for intercept and slopes. The joint distribution of the phenotypic vector **y** is therefore **y** ~ 𝒩 (**X*β*, V**), with **V** = **ZKZ**^*′*^ + **R**, where **R** is a diagonal matrix with heterogeneous residual variances according to field type.

Based on this formulation, the **Mixed Model Equations (MMEs)** for estimating the fixed effects ***β*** and the best linear unbiased predictors (BLUPs) of the random effects **u** are:

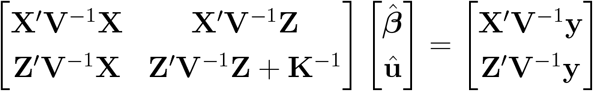

In this implementation, explicit solution of the MMEs was not required, as variance component estimation was performed via restricted maximum likelihood (REML) optimization of the likelihood, and BLUPs were subsequently obtained by solving the conditional expectation equations using the estimated variance components. Nevertheless, the MME formulation provides the theoretical basis underlying the estimation process, ensuring consistency with the standard mixed model framework.

#### 2.2.2 Restricted Maximum Likelihood (REML) Function Implementation

The parameter estimation was performed by maximizing the restricted maximum likelihood (REML). For numerical stability and to ensure that variance components remain within the parameter space, the parameter vector was reparameterized as:

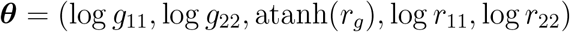

where *g*_11_ and *g*_22_ are the genetic variances for experimental (EXP) and commercial stand (CST) conditions, respectively; *r*_*g*_ is the genetic correlation between field types; and *r*_11_ and *r*_22_ are the residual variances for EXP and CST, respectively. The genetic variance-covariance matrix between field types, **G**_0_, was constructed as:

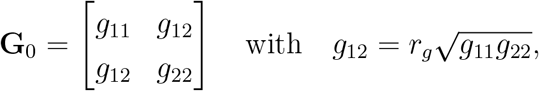

reflecting the assumption of symmetry and consistency between variance and covariance components. To incorporate both the genomic relationship structure and the multiple ECs, I constructed the covariance matrix for the random effects as:

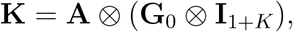

where **A** is the genomic relationship matrix, and **I**_1+*K*_ is an identity matrix representing the random intercept and the *K* = 5 envirotypic covariate slopes. Notably, this product was computationally simplified by first calculating **G**_0_ ⊗ **I**_1+*K*_ to produce a block-diagonal matrix, avoiding the need for an explicit third Kronecker product.

The total variance-covariance matrix of the observations, **V**, was then constructed as:

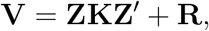

where **Z** is the random effects design matrix, and **R** is a diagonal matrix incorporating heterogeneous residual variances:

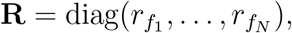

with 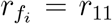 if the *i*-th observation belongs to EXP, or 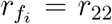 if it belongs to CST. The REML log-likelihood function to be maximized is:

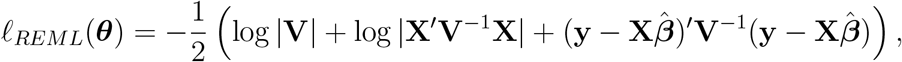

where the fixed effects estimator is given by the generalized least squares (GLS) solution:

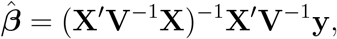

The computational implementation employed **Cholesky decomposition** to efficiently compute **V**^−1^ and log |**V**|, exploiting numerical stability and reducing computational complexity. Specifically, for a matrix **V** with Cholesky factorization **V** = **LL**^*′*^, we have: log |**V**| = 2 ∑_*i*_ log *L*_*ii*_, and the required systems for **V**^−1^**X** and **V**^−1^**y** were solved using forward and backward substitutions, avoiding explicit matrix inversion. The REML function was evaluated in its **negative form**, as minimization is computationally more stable in derivative-free optimization routines. The matrix **V** is large but sparse and structured due to the Kronecker products and the design of **Z**. Residual variances were added element-wise, depending on the field type of each observation.

The function was minimized using the **Bound Optimization BY Quadratic Approximation (BOBYQA)** algorithm, implemented via the `nloptr` package in R (Johnson, 2008), based on the original formulation by Powell (2009). Further details on the parameter initialization and optimization procedure for the R implementation are provided in the following section. BOBYQA is part of the NLopt library, which is also accessible in Python through the `nlopt` package. Additionally, alternative implementations exist, such as `Py-BOBYQA` for Python (Cartis et al., 2019), providing enhanced robustness and flexibility. Other general-purpose derivative-free optimization alternatives include the `COBYLA` algorithm, also available via NLopt, and trust-region-based methods in optimization libraries like `scipy.optimize` (Virtanen et al., 2020).

#### 2.2.3 Parameter Initialization and Optimization Procedure

Initial parameter values were derived empirically from the phenotypic variability observed among genotypes within each field type. Specifically, for each genotype, mean phenotypic values were calculated separately for experimental (EXP) and commercial stand (CST) conditions. These means were then used to compute the phenotypic variances 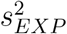 and 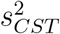, as well as the phenotypic covariance *s*_*EXP,CST*_. The initial value for the genetic correlation parameter was set as the phenotypic correlation:

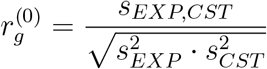

To ensure numerical stability, this value was truncated to lie within (−0.99, 0.99). The initial parameter vector was then defined as:

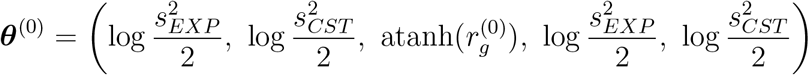

This initialization assumes that both genetic and residual variances for each field type start at similar magnitudes, providing a reasonable starting point for optimization. The optimization was performed using the **BOBYQA** algorithm, as implemented in the `nloptr` package. The bounds for all parameters were set to wide intervals: [−10, 10] for the log-variances and [−5, 5] for the transformed correlation, allowing sufficient flexibility. The relative tolerance for convergence was fixed at 10^−8^, ensuring numerical precision.

The optimization minimized the negative REML log-likelihood function, with convergence diagnostics extracted upon completion, including the estimated REML log-likelihood, the number of iterations, and the total computational time. Specifically, the following diagnostics were recorded:

- **REML log-likelihood:** 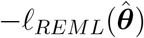.
- **Number of iterations:** as returned by the optimization algorithm.
- **Elapsed computational time:** difference between start and end timestamps.

Upon convergence, the estimated parameter vector 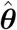 was back-transformed to obtain variance components on the original scale:

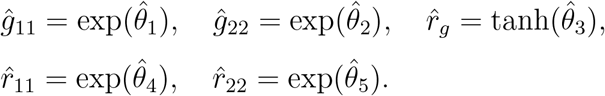

The genetic covariance 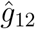 was reconstructed as: 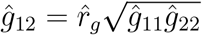 and the genetic variance-covariance matrix **Ĝ**_0_ was obtained accordingly. Similarly, the residual variance-covariance matrix 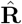 was defined as diagonal with entries 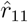 and 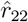.

Following parameter estimation, heritability coefficients were computed for each field type as the ratio between the estimated genetic variance and the total phenotypic variance. Specifically, broad-sense heritability (*H*^2^) was derived under the identity matrix assumption (**A** = **I**), reflecting the total genetic contribution (additive and non-additive) to phenotypic variance, while narrow-sense heritability (*h*^2^) was obtained from the SNP-based genomic relationship matrix, capturing only the additive genetic component. The heritabilities were calculated for experimental (EXP) and commercial stand (CST) conditions as:

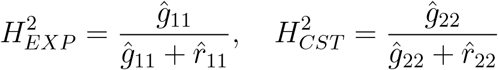

for the identity matrix model, and equivalently for narrow-sense heritabilities 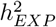 and 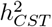 under the genomic model. The difference between *H*^2^ and *h*^2^ was interpreted as an indirect measure of potential non-additive genetic effects not captured by the additive genomic relationship matrix.

### 2.3 Spatial Prediction and Genotype Recommendation Mapping

To assess genotype-specific performance across the environmental space, a spatial prediction procedure was implemented based on the random regression coefficients (BLUPs) for each genotype and field type. The spatial grid for the recommendation maps was defined as a regular lattice from 1 to 10 along both axes, with increments of 0.25 units, resulting in 1369 grid cells (or “pixels”). Since this is a simulation framework, the grid units do not correspond to physical distances (e.g., kilometers or meters).

To synthesize and visualize the main environmental variation across locations, a principal component analysis (PCA) of the standardized ECs was performed. The first two components (PC1 and PC2), jointly explaining a substantial portion of the total environmental variance, were spatially interpolated and mapped. These components provide a reduced-dimensional representation of the dominant environmental gradients potentially influencing genotype performance, facilitating interpretation of the spatial structure of environmental heterogeneity.

Next, genotype-specific predictions were generated by combining the fixed effects (BLUEs), genotype-specific random intercepts, and random slopes associated with each envirotypic covariate. Specifically, for each genotype *g* and field type *f*, the predicted phenotypic value at a given spatial location (*x, y*) was calculated as:

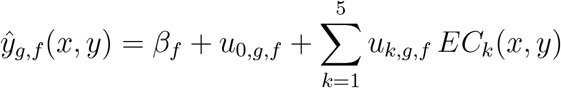

where *β*_*f*_ is the fixed intercept for field type *f, u*_0,*g,f*_ is the genotype-specific random intercept, *u*_*k,g,f*_ are the genotype-specific random slopes, and *EC*_*k*_(*x, y*) are the interpolated envirotypic covariate values at spatial location (*x, y*), with *k* varying from 1 to 5 (ECs), *g* from 1 to 12 (genotypes), and *f* from 1 to 2 (EXP or CST).

The resulting predicted phenotypic values were then compared across genotypes to identify, for each spatial unit, the genotype with the highest predicted performance. This “winner-takes-all” approach allowed the construction of categorical recommendation maps indicating which genotype was predicted to perform best at each spatial location.

The recommendation maps were visualized using faceted plots, separating predictions for experimental (EXP) and commercial stand (CST) scenarios. Distinct colors were assigned to each genotype to facilitate visual discrimination. Notably, the grid-based spatial predictions highlight differential patterns of genotype adaptation, providing a spatially explicit tool to inform genotype deployment strategies under varying environmental conditions.

## 3 Results

Phenotypic variances, based on genotype means across trial locations, were 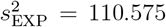 and 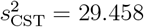, with an observed covariance of *s*_EXP,CST_ = 13.447, resulting in a phenotypic correlation of *r*_p_ = 0.236. These estimates, obtained from *complete cases* (genotypes with phenotypic data in both field types), served as the starting point for the REML estimation.

Figure 1 shows the clustered SNP-based relationship matrix **A**, obtained using Ward’s method. Several groups of genomically related individuals are visually evident: CULF–GEN1– GEN3; CULB–GEN4; GEN2–CULC–CULD; CULA–GEN5; and finally, CULE–CULG. Notably, the pairs GEN5–CULG and CULC–CULE, although not within the same group, exhibit relatively high genomic relationship values. These patterns illustrate both distinct genomic clusters and specific pairwise relationships of interest within the population.

Table 1 presents the REML variance components, heritability estimates, and correlation metrics obtained from the models fitted with both the identity matrix (**A** = **I**_**12**_) and the SNP-based relationship matrix. Noteworthy results include the genetic variances for EXP (*g*_11_ = 36.156 with **A** = **I** and *g*_11_ = 20.294 with SNP-based **A**), as well as the genetic covariance estimates (*g*_12_ = −1.202 for **A** = **I** and *g*_12_ = 1.216 for SNP-based **A**). The genetic correlation was estimated at *r*_*g*_ = −0.130 with **A** = **I**_**12**_ and *r*_*g*_ = 0.218 with SNP-based **A**.

**Table 1.** Comparison of variance and correlation components between the model fitted with **A** = **I**_**12**_ and the SNP-based **A**. The estimates include variance components, correlations, heritability, log-likelihood, and computational diagnostics for each model.

| Parameter | $\mathbf{A} = \mathbf{I}_{12}$ (Identity matrix) | SNP-based $\mathbf{A}$ matrix |
| --- | --- | --- |
| $g_{11}$ (genetic variance EXP) | 36.156 | 20.294 |
| $g_{22}$ (genetic variance CST) | 2.355 | 1.540 |
| $g_{12}$ (genetic covariance) | -1.202 | 1.216 |
| $r_g$ (genetic correlation) | -0.130 | 0.218 |
| $r_{11}$ (residual variance EXP) | 53.726 | 56.898 |
| $r_{22}$ (residual variance CST) | 24.778 | 25.849 |
| $H_{\text{EXP}}^2$ or $h_{\text{EXP}}^2$ (heritability) | 0.402 (broad-sense) | 0.263 (narrow-sense) |
| $H_{\text{EXP}}^2$ or $h_{\text{CST}}^2$ (heritability) | 0.087 (broad-sense) | 0.056 (narrow-sense) |
| $\text{cor}_{\text{EXP}}$ (pred. vs obs.) | 0.886 | 0.869 |
| $\text{cor}_{\text{CST}}$ (pred. vs obs.) | 0.820 | 0.805 |
| $\text{cor}_{\text{pred}}$ (blue+blups EXP-CST) | 0.381 | 0.535 |
| REML log-likelihood | -125.296 | -124.908 |
| Number of iterations | 135 | 141 |
| Elapsed time (seconds) | 0.243 | 0.254 |

*Note:* $g_{11}$ and $g_{22}$ denote genetic variances for experimental (EXP) and commercial stands (CST), respectively. $g_{12}$ represents the genetic covariance between environments. $r_g$ is the genetic correlation. $r_{11}$ and $r_{22}$ are residual variances. $h_{\text{EXP}}^2$ and $h_{\text{CST}}^2$ represent heritability estimates, calculated as $h^2 = g_{ii}/(g_{ii} + r_{ii})$ , where $g_{ii}$ is the genetic variance and $r_{ii}$ is the residual variance for trait $i$ . For the model with $\mathbf{A} = \mathbf{I}_{12}$ , these estimates approximate broad-sense heritability ( $H^2$ ), encompassing total genetic variance. For the SNP-based model, they correspond to narrow-sense heritability ( $h^2$ ), reflecting the proportion of phenotypic variance attributable to additive genetic effects captured by the genomic relationship matrix.

Heritability estimates (*H*^2^ or *h*^2^ = *g*_*ii*_*/*(*g*_*ii*_ + *r*_*ii*_)) were higher with the identity matrix: 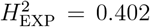 and 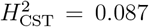. Under the SNP-based model, they decreased to 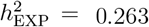 and 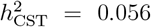, reflecting the narrow-sense additive genetic contributions. The differences—0.139 for EXP and 0.031 for CST—suggest non-additive genetic effects, such as dominance and epistasis, not captured by the additive genomic relationship. Thus, about 13.9% of EXP and 3.1% of CST phenotypic variance may reflect such components.

Residual variances, predictive correlations between observed and predicted values, log-likelihood values, and computational diagnostics are also reported for both models. The SNP-based model exhibited a slightly improved log-likelihood and higher predictive correlation between environments (cor_pred_ = 0.535 vs. 0.381), indicating enhanced predictive consistency when accounting for genomic relatedness.

The fixed intercepts (*β*_0_) and genotype-specific random effects (***u***) estimated for GEN1 to GEN5 under the genomic model reveal systematic differences between experimental (EXP) and commercial stand (CST) conditions. As expected, the fixed intercepts were higher in EXP (*β*_0_ = 247.068) compared to CST (*β*_0_ = 241.896), reflecting distinct baseline performances across field types (Table 2).

**Table 2.** Fixed (*β*_0_) and random (***u***) effects for GEN1 to GEN5 under experimental (EXP) and commercial stand (CST) conditions, including intercepts and slopes for five ECs (EC1 to EC5).

| Gen | Type | $\beta_0$ | $\mathbf{u}_0$ | $\mathbf{u}_1$ | $\mathbf{u}_2$ | $\mathbf{u}_3$ | $\mathbf{u}_4$ | $\mathbf{u}_5$ |
| --- | --- | --- | --- | --- | --- | --- | --- | --- |
| GEN1 | EXP | 247.068 | 4.744 | -3.662 | 3.001 | -3.093 | -3.906 | -1.963 |
|  | CST | 241.896 | 0.786 | -0.783 | 0.516 | -0.389 | -0.599 | -0.477 |
| GEN2 | EXP | 247.068 | -6.113 | 3.928 | -3.953 | 1.894 | 4.795 | 2.656 |
|  | CST | 241.896 | -0.454 | 0.331 | -0.149 | 0.229 | 0.077 | -0.038 |
| GEN3 | EXP | 247.068 | 1.863 | -1.045 | 1.228 | -0.550 | -1.594 | -0.834 |
|  | CST | 241.896 | -0.153 | -0.429 | 0.161 | -0.198 | -0.193 | -0.028 |
| GEN4 | EXP | 247.068 | -0.906 | -0.633 | -0.416 | 1.770 | 0.662 | -0.835 |
|  | CST | 241.896 | -0.532 | 0.190 | -0.135 | 0.101 | 0.147 | 0.213 |
| GEN5 | EXP | 247.068 | -3.254 | 1.099 | -2.085 | 1.838 | 2.659 | 0.764 |
|  | CST | 241.896 | -0.253 | 0.053 | -0.116 | 0.149 | 0.122 | 0.176 |

*Note:* $\beta_0$ represents the fixed intercept (BLUE) for each field type. $\mathbf{u}_0$ is the genotype-specific random intercept (BLUP), and $\mathbf{u}_1$ to $\mathbf{u}_5$ are the random slopes for envirotypic covariates EC1 to EC5, respectively.

The genotype-specific random effects include both the random intercepts (***u***_0_) and the slopes associated with the five ECs (EC1 to EC5). Notably, GEN1 exhibited a positive random intercept in EXP (***u***_0_ = 4.744) and a smaller positive effect in CST (***u***_0_ = 0.786), while GEN2 showed a negative random intercept in EXP (***u***_0_ = −6.113) and a mild negative effect in CST (***u***_0_ = −0.454). Similar patterns were observed for the random slopes, with more pronounced values under EXP conditions for all genotypes, particularly for GEN2 and GEN5, indicating greater genotype-specific sensitivity to ECs in the experimental setting.

From this point forward, results are presented exclusively for the model fitted with the SNP-based relationship matrix (**A**). The model assuming an identity matrix (**A** = **I**_12_) produced a negative genetic covariance between experimental trials (EXP) and commercial stands (CST), with *g*_12_ = −1.202 and a genetic correlation of *r*_*g*_ = −0.130 (Table 1). This estimate suggests a misalignment of genotypic effects across environments, whereby the EXP component disproportionately captured variation, effectively inverting the genetic association between EXP and CST. Given this pattern, analyses and interpretations focus solely on the SNP-based model, which yielded a positive and biologically consistent genetic correlation.

The spatial variation of the ECs and their impact on genotype-specific predictions—where breeding materials evaluated in experiments were predicted across 24 locations as if they were commercial cultivars, using ECs and genomic relationships with the cultivars—is summarized in Figure 4. Panel (A) depicts the spatial interpolation of the first two principal components (PC1 and PC2), which together account for 82.1% of the total environmental variability across the locations. Panel (B) displays predicted phenotypic values for GEN1 to GEN5, obtained under the genomic scenario using the SNP-based matrix; that is, the genotype-specific trajectories represent predicted performances “as cultivar” across all locations.

**Figure 4.**
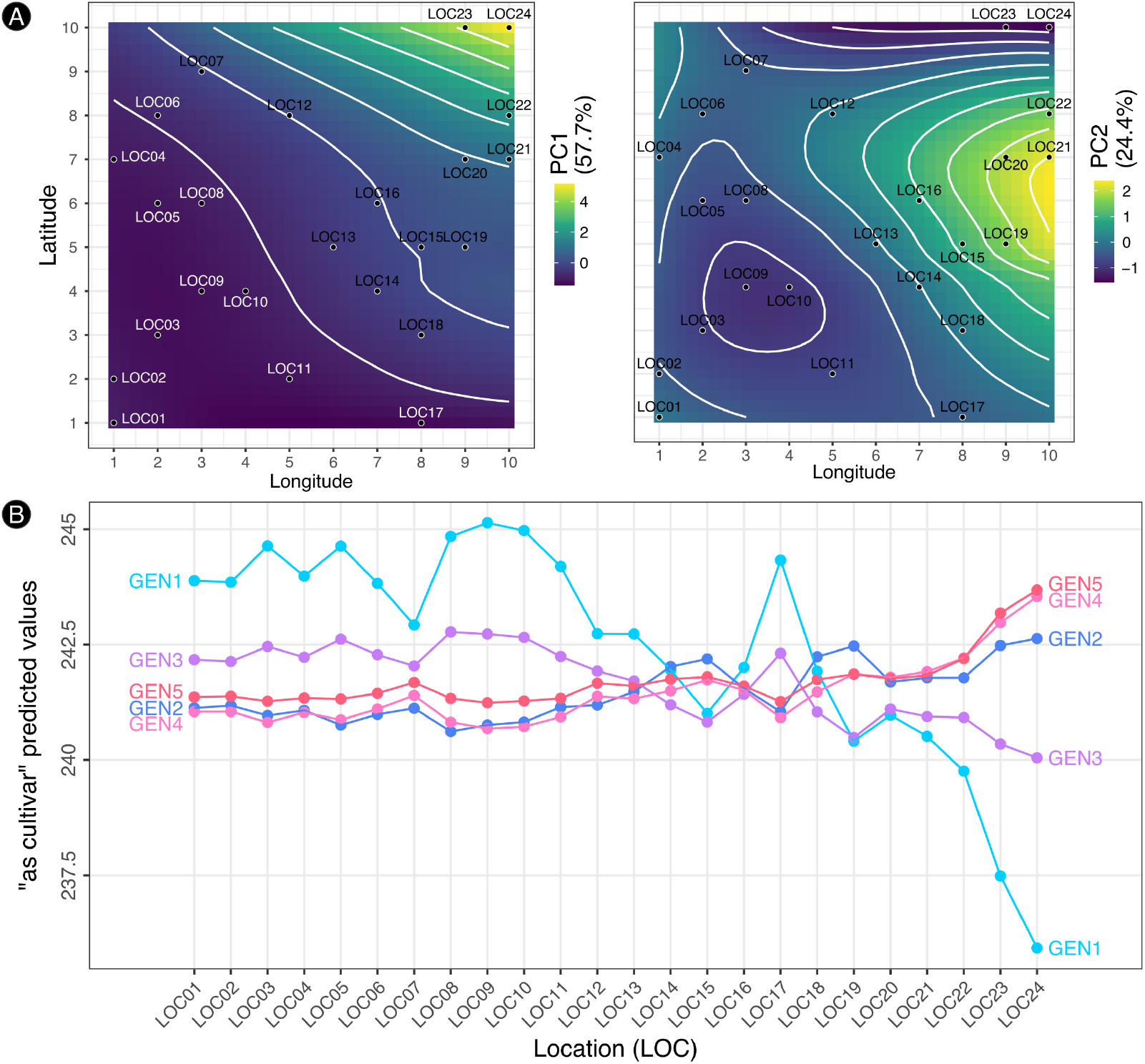
**(A)** Spatial interpolation of the first (PC1, 57.7%) and second (PC2, 24.4%) principal components derived from the EC’s across trial locations (LOC01–LOC24). **(B)** Predicted phenotypic values for the same genotypes using a SNP-based **A** matrix (genomically related genotypes). Lines represent genotype-specific predictions “as cultivar”, with labels at both ends for clarity.

Figure 5 shows the spatial recommendation maps with the top-ranked genotype per spatial unit in both breeding experiments and commercial stand scenarios. The left panel illustrates predicted genotype dominance across experimental environments, while the right reflects their performance as cultivars in commercial stands. GEN5 does not appear in the first panel, as it was not recommended as top-ranked in any spatial unit. Each color marks the genotype with the highest predicted value at each location, highlighting spatial variation in adaptation and supporting environment-specific recommendations.

**Figure 5.**
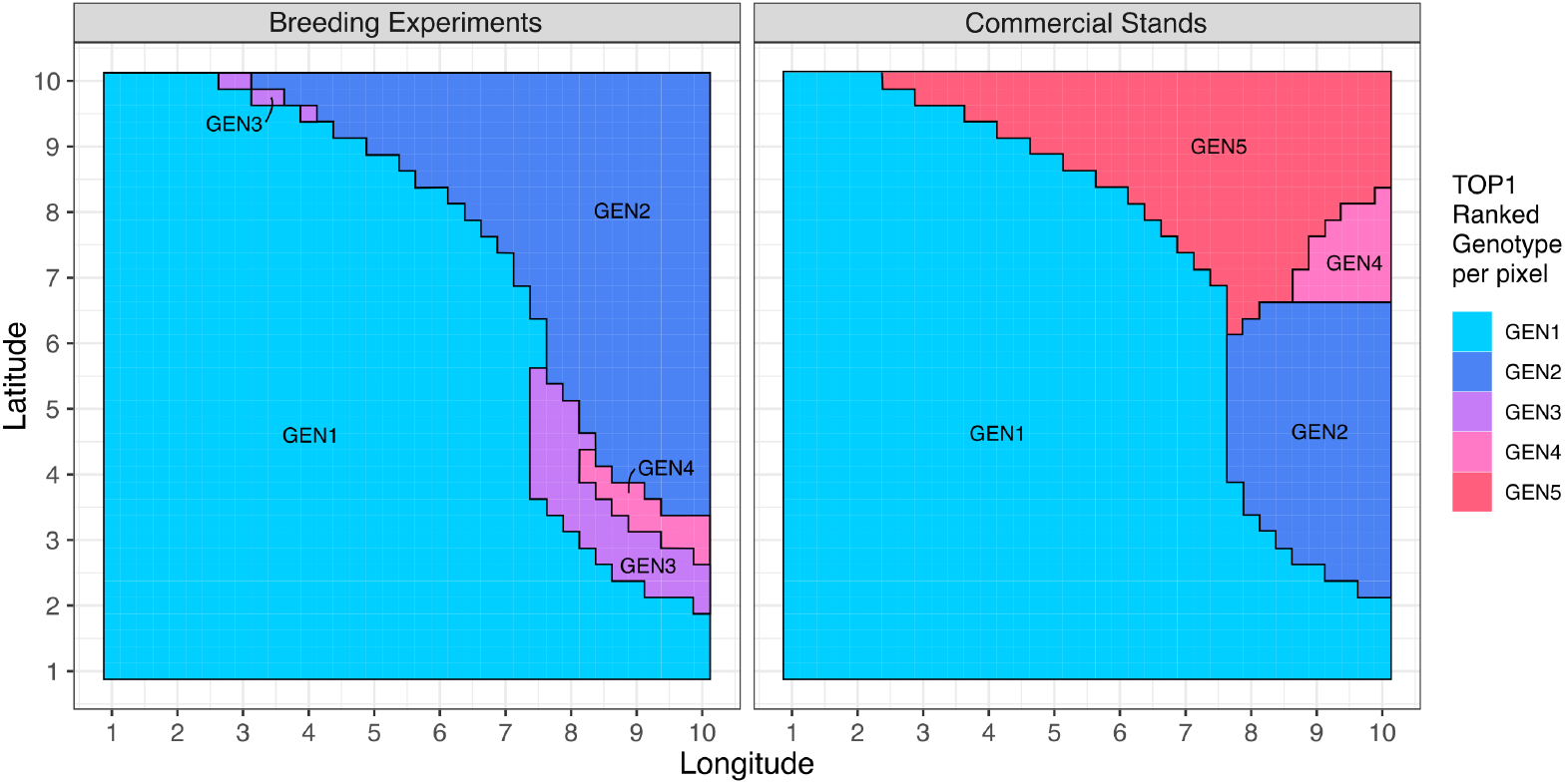
Spatial recommendation map showing the top-ranked genotype per pixel for Breeding Experiments (EXP, figure left) and Commercial Stands (CST, right). Colors represent the genotype achieving the highest predicted value in each spatial unit. This approach supports genotype-specific recommendations based on environmental gradients and genomic SNP-based information.

## 4 Discussion

### 4.1 Estimates from the REML Model: Methodological Reflections and Modeling Experience

The choice of the relationship matrix structure is fundamental in mixed models, directly influencing the estimation of variance components and subsequent predictions (Kelly et al., 2009; Amadeu et al., 2023). The identity matrix is often employed as a baseline in situations where no explicit pedigree or genomic information is available. However, such simplification assumes unrelated genotypes and disregards the realized genetic similarities captured by genomic markers. In this study, the comparison between using **A** = **I**_**12**_ and the SNP-based genomic relationship matrix highlighted the role that incorporating genomic information plays in obtaining biologically coherent estimates and reliable predictions. This observation is consistent with previous findings demonstrating that integrating genomic data early in the breeding pipeline enhances predictive accuracy and reduces phenotyping costs (Beyene et al., 2021; de Sousa et al., 2021).

The comparative results demonstrated that, when using **A** = **I**_**12**_, the estimated genetic covariance (*g*_12_ = −1.202) and genetic correlation (*r*_*g*_ = −0.130) between experimental trials (EXP) and commercial stands (CST) were negative, indicating an implausible inversion of genetic effects across these environments. In contrast, fitting the model with the SNP-based matrix produced a positive genetic covariance (*g*_12_ = 1.216) and correlation (*r*_*g*_ = 0.218), consistent with expectations for shared genetic control across environments. Such improvements echo prior observations that explicit modeling of genotype-by-environment interactions mitigates counterintuitive or unstable predictions, particularly under scenarios of incomplete trials or extrapolative inference (Wittenburg et al., 2016; Roorkiwal et al., 2018).

Methodologically, this experience underscores the risks of using **A** = **I**_**12**_ in datasets with disconnected genotype-environment combinations, a common occurrence in breeding programs where not all genotypes are tested across all environments (Hartung & Piepho, 2021; Roorkiwal et al., 2018). In this study, the data exhibited spatial and structural imbalance, notably with an accumulation of experimental observations concentrated in the southwestern region of the study area. Such data unbalancing exacerbates the risk of spurious estimates when using simplistic assumptions of genetic independence. Indeed, prior studies have shown that modeling genotype-specific reaction norms or leveraging environmental genomic selection frameworks can mitigate the risks associated with extrapolative errors in similar contexts (Ly et al., 2018; Costa-Neto et al., 2021b, Costa-Neto et al., 2023; Gevartosky et al., 2023; Halpin-McCormick et al., 2025). Incorporating genomic information via the SNP-based matrix introduced essential regularization, mitigating artifacts such as the observed “flip” in genetic effects between EXP and CST field types, and providing more stable estimates for variance components and genetic correlations.

From a computational perspective, the REML approach implemented here proved reasonably efficient for the dataset analyzed, successfully estimating an unstructured (also called ‘UN’ or ‘UNS’ by some available mixed model packages/programs) genetic variance-covariance matrix in a bivariate model with ECs. Nonetheless, further optimization is warranted for scaling the approach to larger datasets or more complex models. Potential improvements include exploiting the inherent sparsity of the random effects design matrix **Z**, utilizing more efficient algorithms for Kronecker product computations, or adopting alternative optimization strategies such as those integrating stochastic approximation methods (Barreto et al., 2024). These considerations are particularly pertinent given that multivariate or machine learning approaches, while enhancing predictive accuracy, can impose additional computational burdens requiring careful algorithmic optimization (Gill et al., 2021; Xu et al., 2022; Mora-Poblete et al., 2023; Crossa et al., 2024).

The present framework could be extended to incorporate additional phases of the breeding program beyond the bivariate specification. Although referring to the same trait, its expression across phases—early trials, advanced tests, and commercial deployment (farming)—constitutes distinct yet genetically correlated traits. A multi-phase model would leverage genetic covariances among these stages (Bhosale et al., 2025), computed through genotypes common across phases, such as selected lines, relatives, or standard checks. These shared genotypes provide links for estimating covariances, enabling more informed predictions (Auinger et al., 2016). While such extensions increase computational complexity, the structured design of the random effects matrix **Z** and efficient covariance representations offer a practical scaling path. Moreover, the approach remains compatible with paradigms integrating continuous environmental gradients within broader reaction norm frameworks (Ly et al., 2018; Resende et al., 2024).

Complementing the mixed model strategy with machine learning (ML) and deep learning methods could be considered. These approaches have shown promise in capturing complex genotype-by-environment interactions, particularly when dealing with large-scale, multi-trait data (Mora-Poblete et al., 2023; Crossa et al., 2024). However, it is important to recognize that model interpretability and computational efficiency often need to be balanced against predictive gains. The use of derivative-free REML (DF-REML) approaches, as implemented here, opens avenues for modern computational strategies, including parallelization on GPUs or NPUs (Graphics and Neural Processing Unit, respectively), and potential integration within machine learning architectures (Xu et al., 2022). In this sense, mixed models remain quietly elegant—mathematical constructs that, despite their traditional roots, continue to serve as the computational backbone of sophisticated predictive frameworks. As emphasized in recent reviews, hybrid strategies combining statistical models with ML algorithms may offer a middle ground, leveraging the strengths of both paradigms for genomic prediction in diverse breeding scenarios (Crossa et al., 2024).

### 4.2 Predictive performance and spatial recommendations for breeding genotypes

Breeding (candidate) genotypes (GEN’s) were evaluated exclusively in experimental trials (LOC01 to LOC05), whereas commercial cultivars (CUL’s) were assessed in on-farm commercial stands (LOC06 to LOC24). Some CUL’s were also used as checks in the experiments, a common practice for comparative purposes or as benchmarks to be surpassed (Eskridge and Mumm, 1992; Crossa et al., 2021). It’s also important to notice that, in many cases, a cultivar released was previously an experimental/candidate genotype, reflecting the dynamism of breeding programs. Thus, the distinction between experimental genotypes and commercial cultivars can, in part, represent the temporal phase of evaluation, reinforcing the need to consider the complete history of materials when interpreting performance and genomic relationship data.

Regarding genomic structure, CULF, GEN1, and GEN3 form a genetically close group, with GEN1 showing the best phenotypic performance, while GEN3, evaluated in only two locations, showed stable and satisfactory performance (Figure 1). Similarly, CULA and GEN5 exhibit high genomic similarity; moreover, GEN5 is also closely related to the cultivar CULG, which is notably deployed in the northeastern region of the study area. Consistently, the genomic model recommended GEN5 for that region, as shown in Figure 5 (right panel). However, GEN5 was assessed only in one environment (LOC01), with low yield, limiting robust inferences about its adaptability. These cases illustrate that genomic proximity alone is insufficient to guarantee superior phenotypic performance or broad recommendation, emphasizing the need to account for genotype-by-environment interaction in breeding material evaluation (Bustos-Korts et al., 2016; Costa-Neto et al., 2021b, Costa-Neto et al., 2023).

On the other hand, CULB and GEN4 stand out for their genomic proximity and for both being evaluated in the experimental trials, where they exhibited similar phenotypic behaviors. However, CULB showed poor performance in commercial stands, suggesting that in environments where CULB does not adapt well, GEN4 is also unlikely to be recommended—an inference supported by the spatial recommendation maps (Figures 4 and 5). The group CULC, CULD, and GEN2 appears closely clustered in the dendrogram; CULC and CULD are excellent commercial materials, while GEN2, although genetically similar, had the worst performance in experiments. Nevertheless, the fitted model indicated that GEN2 would be recommended as the best genotype in specific locations; Bustos-Korts et al. (2016) reinforce the value of predictive models beyond raw phenotypic evaluation.

The experimental trials (LOC01–LOC05) are concentrated exclusively in the southwestern region of the study area, where environmental conditions are relatively homogeneous. This limited spatial distribution compromises the representativeness of the experimental data for the entire area, especially regarding the central and northeastern regions, which present environments with distinct climatic and edaphic characteristics. Therefore, relying solely on observed experimental data for recommendations would be inadequate, highlighting the importance of predictive approaches that incorporate ECs and genomic relationship structures to extend inferences to environments not directly tested (Costa-Neto et al., 2023). Identifying genotype-specific adaptation across heterogeneous environments can improve breeding outcomes, given that targeted selection achieves greater genetic gains than wide adaptation (Annicchiarico, 2021; Cruz et al., 2025).

The predictive analysis conducted in this study allowed simulation of the in-field performance of the breeding genotypes across all locations, as if they were commercial cultivars, using information on genomic relationships and multi-environmental gradients. This procedure evidenced, for instance, that genotypes such as GEN4 and GEN5, which showed modest or limited performance in the experimental trials, demonstrated specific adaptive potential for certain regions, especially in the northeastern portion of the study area. Thus, predictive modeling revealed patterns and opportunities that would not be perceptible based solely on the phenotypic data collected in the experiments. Accurate variance component estimation and predictive inference can still be achieved in multi-environment trials despite data imbalance or missing observations, as demonstrated in simulation studies (Hartung & Piepho, 2021).

Although this study demonstrated the potential of predictive extrapolation across environments, spatial recommendation maps can guide genotype selection in specific regions, reducing the need for extensive multi-environment trials (METs), optimizing resources, and accelerating breeding (Annicchiarico, 2021; Resende et al., 2021). In the maps (Figure 5), only the top-ranked genotype per pixel is shown, but others should be considered. For example, a 2^nd^ or 3^rd^ ranked genotype may be preferred if offering logistical advantages or resistance to local stresses (e.g., frost, nematodes, etc.), critical in genotypic planting recommendation decisions.

The current model uses only five ECs, which is limited in the Enviromics context and could be improved by incorporating additional covariates (Costa-Neto et al., 2021a), as well as through engineered enviromic markers and ensemble modeling strategies, as proposed by Resende et al. (2021), potentially increasing predictive accuracy. Additionally, this study lacks formal validation procedures, such as cross-validation or independent datasets, whose future implementation will reinforce the robustness and practical applicability of the model. Nevertheless, it serves as a methodological protocol, focusing on conceptual demonstration and technical implementation, constituting a solid foundation for future, more robust applications.

## Notes

### Competing Interest Statement

The authors have declared no competing interest.

## References

Amadeu, R. R., Garcia, A. A. F., Munoz, P. R., & Ferrão, L. F. V. (2023). AGHmatrix: genetic relationship matrices in R. Bioinformatics, 39(7), btad445.

Annicchiarico, P. (2021). Breeding gain from exploitation of regional adaptation: An alfalfa case study. Crop Science, 61(4), 2254–2271.

Aparicio, J., Gezan, S. A., Ariza-Suarez, D., Raatz, B., Diaz, S., Heilman-Morales, A., & Lobaton, J. (2024). Mr. Bean: a comprehensive statistical and visualization application for modeling agricultural field trials data. Frontiers in Plant Science, 14, 1290078.

Auinger, H.-J., Schönleben, M., Lehermeier, C., Schmidt, M., Korzun, V., Geiger, H. H., et al. (2016). Model training across multiple breeding cycles significantly improves genomic prediction accuracy in rye (Secale cereale L.). Theoretical and Applied Genetics, 129(11), 2043–2053.

Barreto, C. A. V., Dias, K. O. G., de Sousa, I. C., Azevedo, C. F., Nascimento, A. C. C., Guimarães, L. J. M., et al. (2024). Genomic prediction in multi-environment trials in maize using statistical and machine learning methods. Scientific Reports, 14, 1062.

Bates, D., Mächler, M., Bolker, B., & Walker, S. (2015). Fitting linear mixed-effects models using lme4. Journal of Statistical Software, 67(1), 1–48.

Bernardo, R. (2010). Breeding for quantitative traits in plants (2nd ed.). Woodbury, MN: Stemma Press.

Beyene, Y., Semagn, K., et al. (2021). Application of genomic selection at the early stage of breeding pipeline in tropical maize. Frontiers in Plant Science, 12, 685488.

Bhosale, S. D., et al. (2025). OneRice breeding framework: An end-to-end system to develop better varieties faster. Crop Science, 65(1), 123–135.

Bustos-Korts, D., Malosetti, M., Chapman, S., & van Eeuwijk, F. (2016). Modelling of genotype by environment interaction and prediction of complex traits across multiple environments as a synthesis of crop growth modelling, genetics and statistics. In X. Yin & P.C. Struik (Eds.), Crop Systems Biology: Narrowing the gaps between crop modelling and genetics (pp. 55–82). Cham: Springer International Publishing.

Cartis, C., Fiala, J., Marteau, B., & Roberts, L. (2019). Improving the flexibility and robustness of model-based derivative-free optimization solvers. ACM Transactions on Mathematical Software, 45(3), 32:1–32:41.

Cooper, M., & Messina, C. D. (2021). Can we harness “Enviromics” to accelerate crop improvement by integrating breeding and agronomy? Frontiers in Plant Science, 12, 735143.

Costa-Neto, G., Galli, G., Carvalho, H. F., Crossa, J., & Fritsche-Neto, R. (2021). EnvR-type: a software to interplay enviromics and quantitative genomics in agriculture. G3: Genes—Genomes—Genetics, 11(4), jkab040.

Costa-Neto, G., Crossa, J., & Fritsche-Neto, R. (2021). Enviromic assembly increases accuracy and reduces costs of the genomic prediction for yield plasticity in maize. Frontiers in Plant Science, 12, 717552.

Costa-Neto, G., Crespo-Herrera, L., Fradgley, N., Gardner, K., Bentley, A. R., Dreisigacker, S., & Crossa, J. (2023). Envirome-wide associations enhance multi-year genome-based prediction of historical wheat breeding data. G3, 13(2), jkac313.

Crossa, J., Perez-Rodriguez, P., Cuevas, J., Montesinos-López, O., Jarquín, D., de Los Campos, G., et al. (2021). Genomic selection in plant breeding: Methods, models, and perspectives. Trends in Plant Science, 26(10), 975–996.

Crossa, J., Martini, J. W. R., Vitale, P., Pérez-Rodríguez, P., Costa-Neto, G., Fritsche-Neto, R., et al. (2024). Expanding genomic prediction in plant breeding: harnessing big data, machine learning, and advanced software. Trends in Plant Science.

Cruz, D. D., Heinemann, A. B., Marcatti, G. E., & Resende, R. T. (2025). Defining the target population of environments (TPE) for enviromics studies using R-based GIS tools. Crop Breeding and Applied Biotechnology, 25(1), e50822519.

de Sousa, K., van Etten, J., Poland, J., Fadda, C., Jannink, J.-L., Kidane, Y. G., et al. (2021). Data-driven decentralized breeding increases prediction accuracy in a challenging crop production environment. Communications Biology, 4, 944.

Eskridge, K. M., & Mumm, R. F. (1992). Choosing plant cultivars based on the probability of outperforming a check. Theoretical and Applied Genetics, 84, 494–500.

Gevartosky, R., Carvalho, H. F., Costa-Neto, G., Montesinos-López, O. A., Crossa, J., & Fritsche-Neto, R. (2023). Enviromic-based kernels may optimize resource allocation with multi-trait multi-environment genomic prediction for tropical maize. BMC Plant Biology, 23(1), 10.

Gill, H. S., Halder, J., Zhang, J., Brar, N. K., Rai, T. S., Hall, C., et al. (2021). Multitrait multi-environment genomic prediction of agronomic traits in advanced breeding lines of winter wheat. Frontiers in Plant Science, 12, 709545.

Halpin-McCormick, A., Campbell, Q., Negrão, S., Morrell, P. L., Hübner, S., Neyhart, J. L., & Kantar, M. B. (2025). Environmental genomic selection to leverage polygenic local adaptation in barley landraces. Communications Biology, 8(1), 618.

Hartung, J., & Piepho, H. P. (2021). Effect of missing values in multi-environmental trials on variance component estimates. Crop Science, 61(6), 4087–4097.

Heslot, N., Akdemir, D., Sorrells, M. E., & Jannink, J.-L. (2014). Integrating environmental covariates and crop modeling into the genomic selection framework to predict genotype-by-environment interactions. Theoretical and Applied Genetics, 127(2), 463–480.

Heslot, N., Jannink, J.-L., & Sorrells, M. E. (2015). Perspectives for genomic selection applications and research in plants. Crop Science, 55(1), 1–12.

Jarquín, D., Crossa, J., Lacaze, X., Du Cheyron, P., Daucourt, J., Lorgeou, J., et al. (2014). A reaction norm model for genomic selection using high-dimensional genomic and environmental data. Theoretical and Applied Genetics, 127(3), 595–607.

Johnson, S. G. (2008). The NLopt nonlinear-optimization package. https://github.com/stevengj/nlopt.

Kelly, A. M., Cullis, B. R., Gilmour, A. R., Eccleston, J. A., & Thompson, R. (2009). Estimation in a multiplicative mixed model involving a genetic relationship matrix. Genetics Selection Evolution, 41(1), 33.

Laidig, F., Drobek, T., & Meyer, U. (2008). Genotypic and environmental variability of yield for cultivars from 30 different crops in German official variety trials. Plant Breeding, 127(6), 541–547.

Lopez-Cruz, M. A., Crossa, J., Bonnett, D., Dreisigacker, S., Poland, J., Jannink, J.-L., & Pérez-Rodríguez, P. (2020). Increased prediction accuracy in wheat breeding trials using a marker × environment interaction genomic selection model. G3: Genes, Genomes, Genetics, 10(7), 2625–2637.

Ly, D., Huet, S., Gauffreteau, A., Rincent, R., Touzy, G., Mini, A., et al. (2018). Whole-genome prediction of reaction norms to environmental stress in bread wheat (Triticum aestivum L.) by genomic random regression. Field Crops Research, 216, 32–41.

Messina, C. D., Technow, F., Tang, T., Totir, L. R., Gho, C., & Cooper, M. (2018). Leveraging biological insight and environmental variation to improve phenotypic prediction: Integrating crop growth models (CGM) with whole genome prediction (WGP). European Journal of Agronomy, 100, 151–162.

Milad, M., Cooper, M., Messina, C. D., & Chapman, S. (2023). Advances in digital phenotyping and enviromic modeling for plant breeding under climate change. Plant Phenomics, 5, 1234567.

Monteverde, E., Resende, M. F. R., Ferrão, L. F. V., & Munoz, P. R. (2023). Multi-trait multi-environment genomic prediction models using enviromic data to improve prediction accuracy in forest trees. G3: Genes, Genomes, Genetics, 13(2), jkac321.

Mora-Poblete, F., Maldonado, C., Henrique, L., Uhdre, R., Scapim, C. A., Mangolim, C. A., et al. (2023). Multi-trait and multi-environment genomic prediction for flowering traits in maize: a deep learning approach. Frontiers in Plant Science, 14, 1153040.

Podlich, D. W., Cooper, M., & Basford, K. E. (2001). Modeling plant breeding programs as search strategies on a complex response surface. Crop Science, 41(3), 624–639.

Powell, M. J. D. (2009). The BOBYQA algorithm for bound constrained optimization without derivatives. Technical Report NA2009/06, Department of Applied Mathematics and Theoretical Physics, University of Cambridge.

Resende, R. T., Piepho, H.-P., Rosa, G. J. M., Silva-Junior, O. B., Silva, F. F. E., de Resende, M. D. V., & Grattapaglia, D. (2021). Enviromics in breeding: applications and perspectives on envirotypic-assisted selection. Theoretical and Applied Genetics, 134(1), 95–112.

Resende, R. T., Hickey, L., Amaral, C. H., Peixoto, L. L., Marcatti, G. E., & Xu, Y. (2024). Satellite-enabled enviromics to enhance crop improvement. Molecular Plant, 17(6), 848–866.

Resende, R. T., Xavier, A., Silva, P. I. T., Resende, M. P., Jarquin, D., & Marcatti, G. E. (2025). GIS-based G× E modeling of maize hybrids through enviromic markers engineering. New Phytologist, 245(1), 102–116.

Roorkiwal, M., Jarquin, D., Singh, M., Gaur, P. M., Bharadwaj, C., Rathore, A., et al. (2018). Genomic-enabled prediction models using multi-environment trials to estimate the effect of genotype×environment interaction on prediction accuracy in chickpea. Scientific Reports, 8, 11701.

Tilhou, N. W., & Casler, M. D. (2022). Genetic correlations between switchgrass performance in sward conditions and surrogate measures. Crop Science, 62(4), 1511–1521.

Trevisan, B. A., Junqueira, V. S., Florencio, B. D. M., Coelho, A. S., Marcatti, G. E., & Resende, R. T. (2025). A framework for building enviromics matrices in mixed models. arXiv preprint 2501.04147.

Virtanen, P., Gommers, R., Oliphant, T. E., et al. (2020). SciPy 1.0: fundamental algorithms for scientific computing in Python. Nature Methods, 17, 261–272.

Wittenburg, D., Teuscher, F., Klosa, J., & Reinsch, N. (2016). Covariance between genotypic effects and its use for genomic inference in half-sib families. G3: Genes, Genomes, Genetics, 6(9), 2761–2772.

Xu, Y., Zhang, X., Li, H., Zheng, H., Zhang, J., Olsen, M. S., Varshney, R. K., Prasanna, B. M., & Qian, Q. (2022). Smart breeding driven by big data, artificial intelligence, and integrated genomic-enviromic prediction. Molecular Plant, 15(11), 1664–1695.

